## Supplementary Information for "Expanding the pool of public controls for GWAS via a method for combining genotypes from arrays and sequencing"

### Supplementary Figures

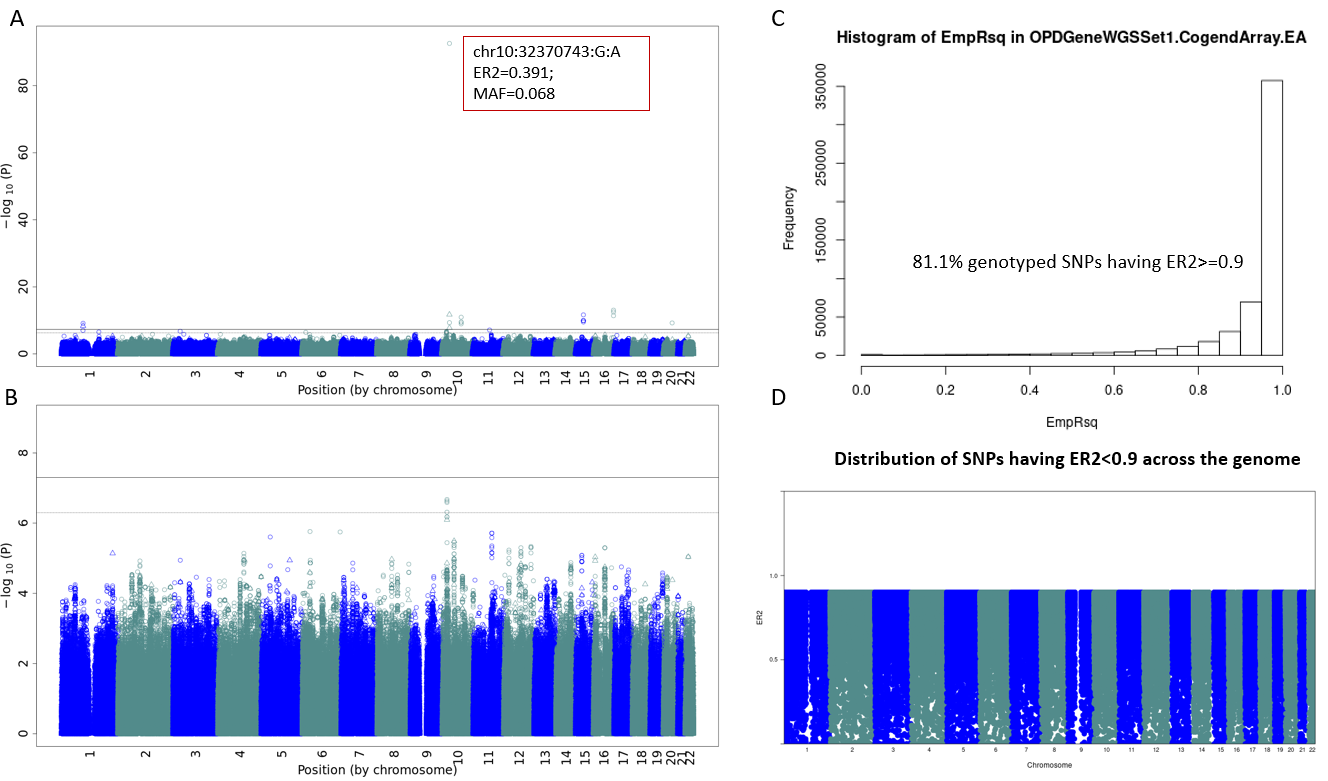

**Supplementary Figure 1: Utility of the empirical R^2^ (ER^2^) filter**. Taking the test between COGEND and COPDGene EA sample comparison (Analysis 1 in the control of false positive tests) as an example, we conducted GWAS with COGEND EA array data as cases and COPDGene EA1 WGS data as controls. (A) Before applying the ER^2^ filter, many false positives resulted due to the poor ER^2^. (B) With the ER^2^ filter of 0.9, the false positives were well controlled. (C) The histogram shows the distribution of ER^2^ in all genotyped SNPs. (D) The scatter plot of the outlier SNPs with ${ER}^{2}<0.9$ were evenly distributed across the genome. The Y-axis is the value of ${ER}^{2}$.

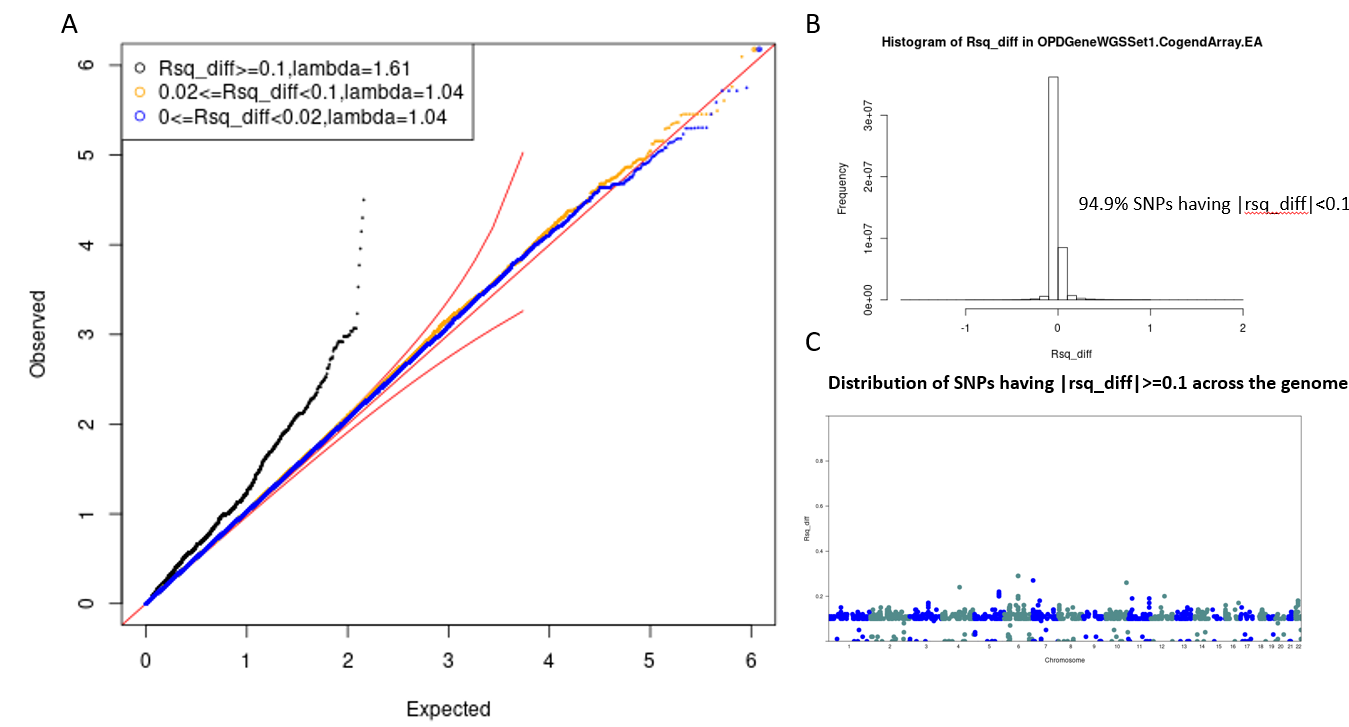

**Supplementary Figure 2: Utility of R^2^ Difference Filter**. Taking the test between COGEND and COPDGene EA sample comparison (Analysis 1 in the control of false positive tests) as an example, we conducted GWAS with COGEND EA array data as cases and COPDGene EA1 WGS data as controls. (A) The QQ plot shows the P-values for the subset of SNPs with $\left| R_{array}^{2}-R_{WGS}^{2} \right|\geq0.1$ has problematic inflation (black dots), but good for other SNPs (orange and blue dots). (B) The histogram shows the distribution of Rsq_diff ($\left| R_{array}^{2}-R_{WGS}^{2} \right|)$ in all SNPs. (C) The scatter plot of the outlier SNPs with $\left| R_{array}^{2}-R_{WGS}^{2} \right|\geq0.1$ are evenly distributed across the genome. The Y-axis is the value of Rsq_diff.

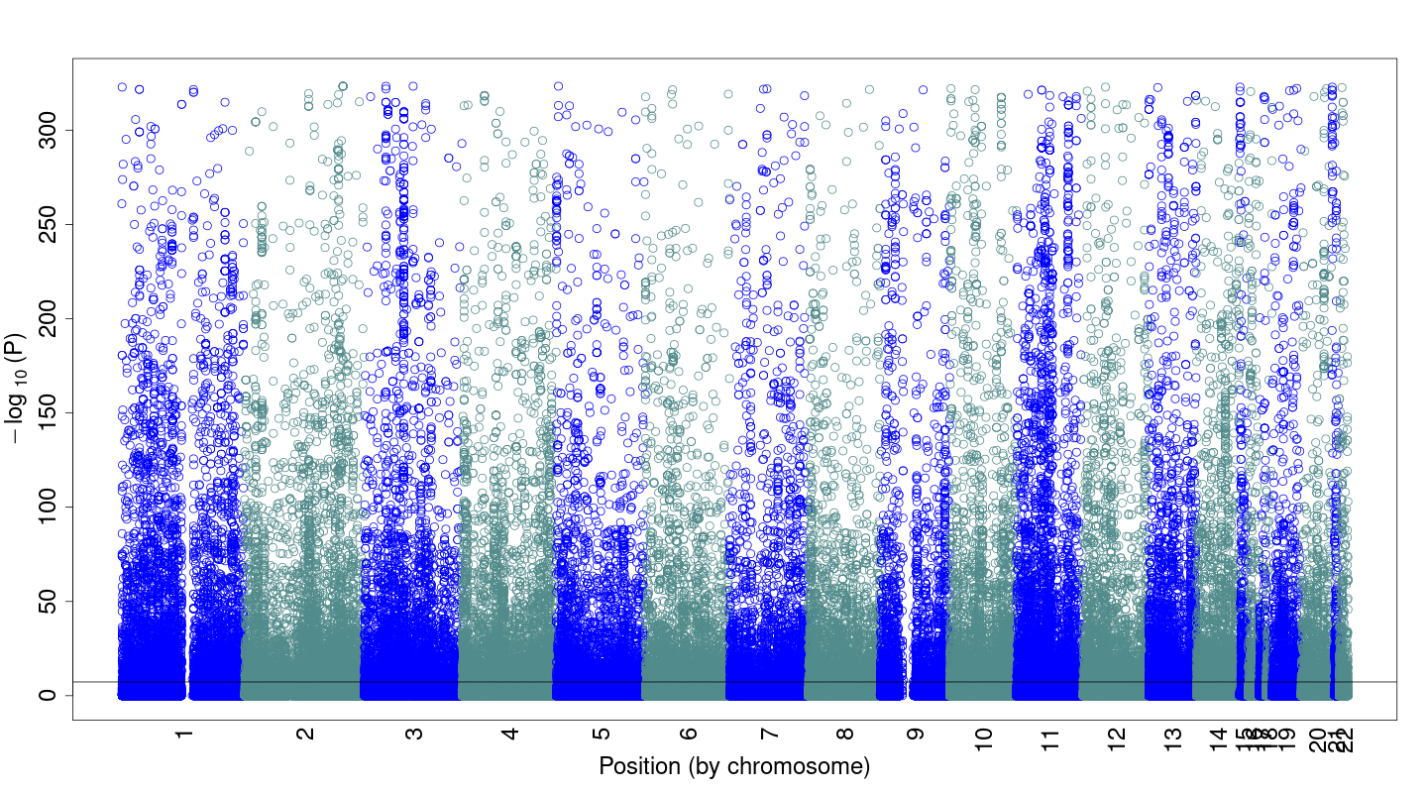

**Supplementary Figure 3: False Positives with TOPMed phased data.** Manhattan plot comparing WGS phased vs array data from the same set of COPDGene samples. The WGS data is phased by TOPMed based on all genotypes on all samples within the program. Therefore, the array and WGS data have been phased with different single nucleotide polymorphisms and sample set. We believe the difference in phasing is the cause of the false positives and these are corrected in the results shown in Supplementary Figure 4.

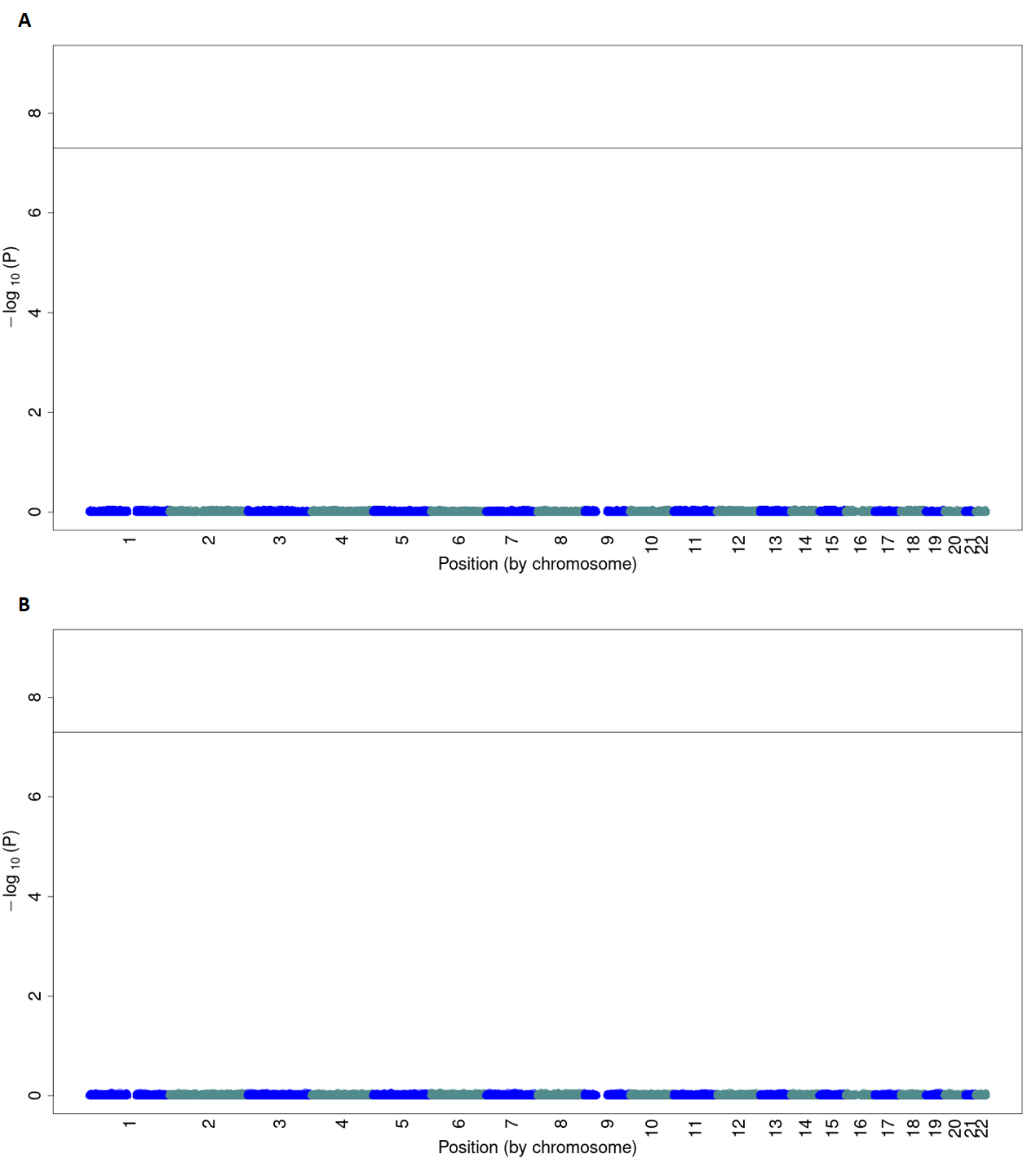

**Supplementary Figure 4: Technical reproducibility across genotyping technology.** Manhattan plots comparing WGS vs. array genotyping data from the same set of COPDGene samples of (A) European ancestry (N=6,501) and (B) African American ancestries (N=3,235) after applying GAWMerge.

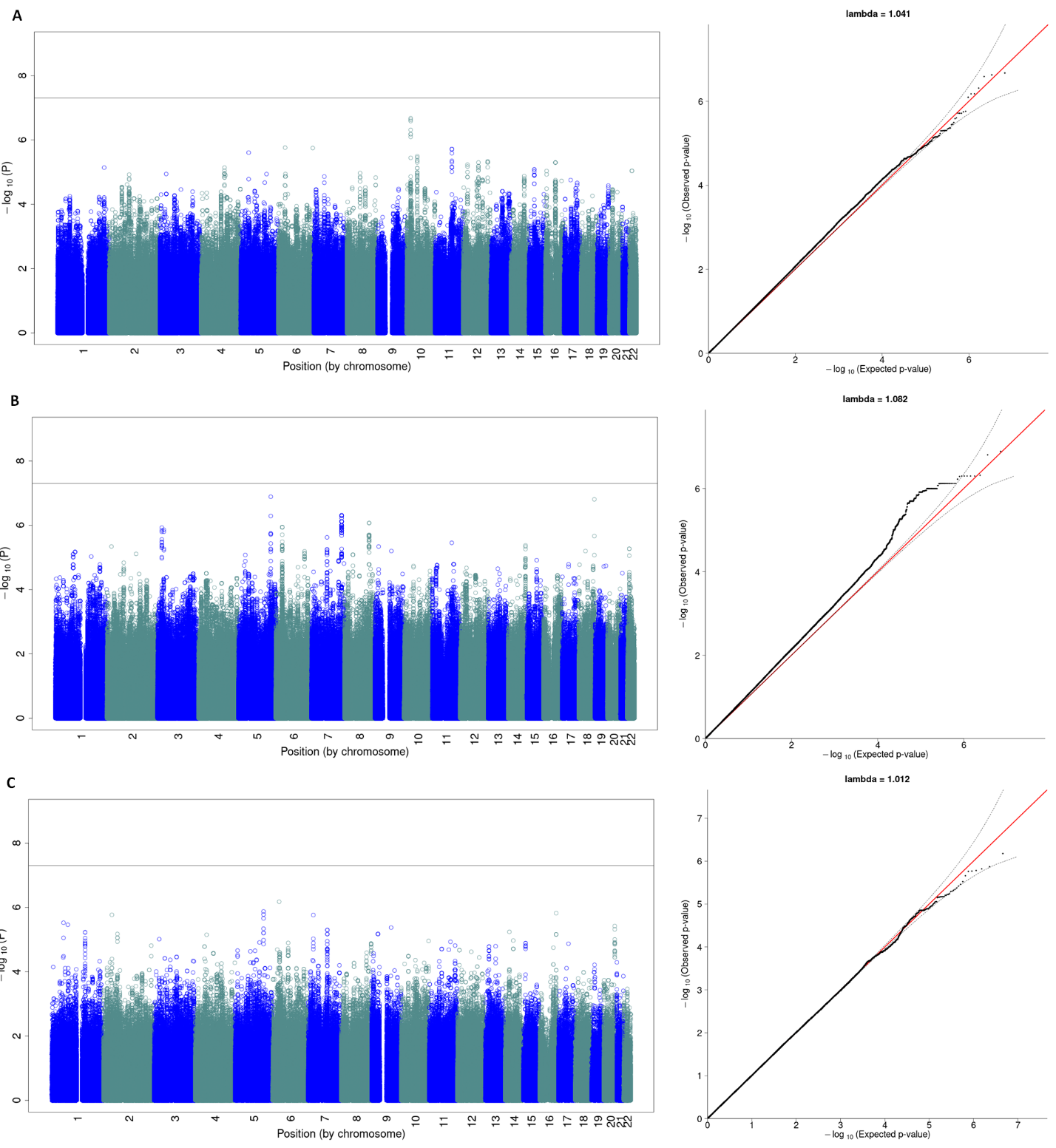

**Supplementary Figure 5: Controlling type 1 error across genotype technology.** Manhattan and quantile-quantile (QQ) plots for the analyses to control type I error for the (A) COGEND EA array data (N=1,961) vs. COPDGene EA1 WGS data (N=3,251), (B) COPDGene EA2 array data (N=3,251) vs. ECLIPSE EA WGS data (N=1,461), and (C) COGEND AA array data (N=712) Vs. COPDGene AA WGS data (N=1,710) analyses. As all of the cohorts, COGEND, COPDGene, and ECLIPSE are smoking cohorts along with the COPD case distributed across both classes being tested, we see no genome-wide signal, thus controlling type 1 error.

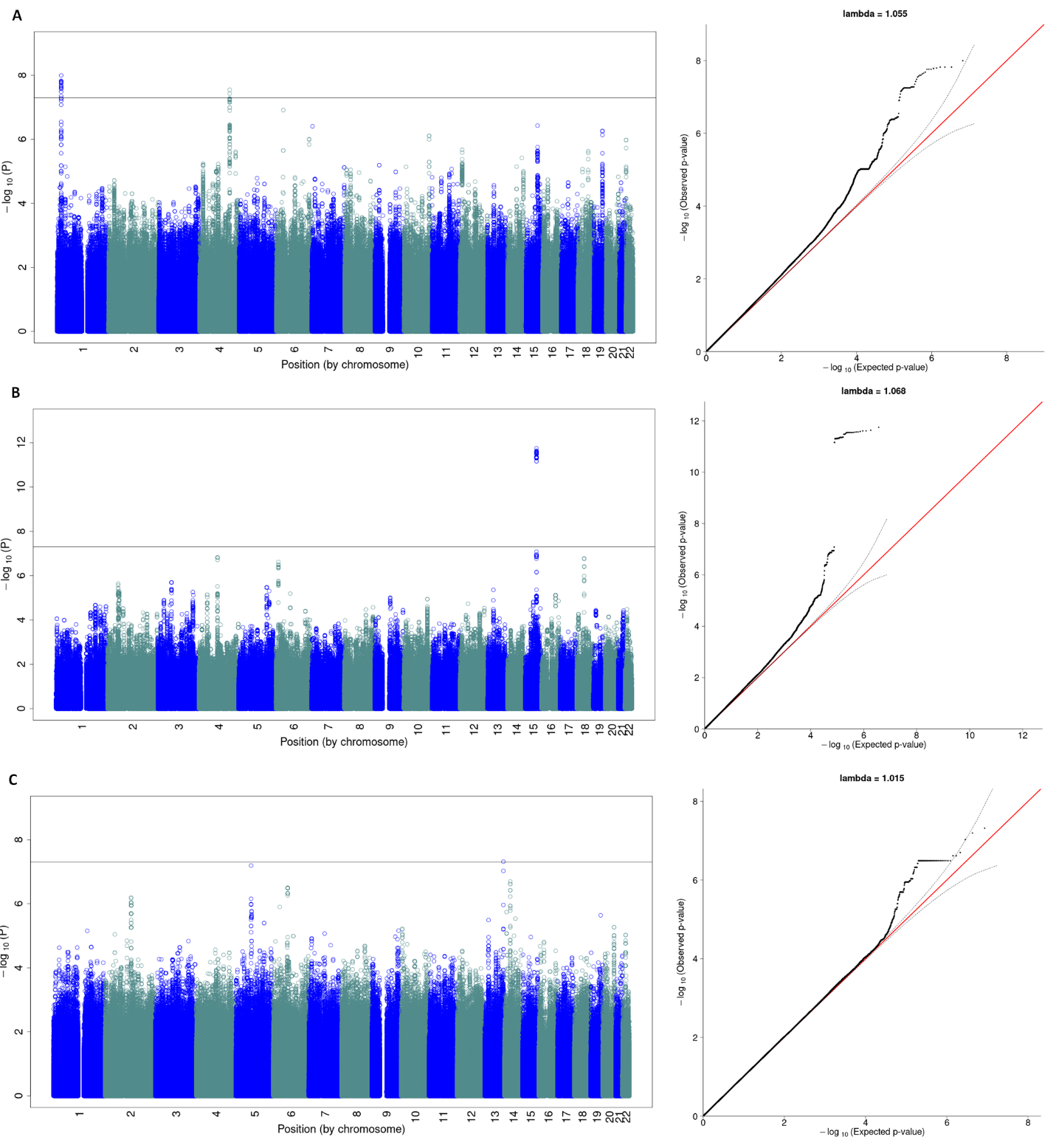

**Supplementary Figure 6: Recovery of known loci across technology.** Manhattan and quantile-quantile (QQ) plots for the analyses to recover COPD GWAS signals for the (A) COGEND EA array data as smoking controls (N=1,961) and COPDGene EA with COPD WGS data as cases (N=2,736), (B) ECLIPSE EA with COPD array data as cases (N=1,764) and COPDGene EA with no COPD WGS data as smoking controls (N=2,475), and (C) COGEND AA array data as smoking controls (N=712) and COPDGene AA with COPD WGS data as cases (N=813) analyses.

### Supplementary Tables

**Supplemental Table 1:** TOPMed Consortium Banner Authors

| **Name** | **Institution(s)** |
| --- | --- |
| Abe, Namiko | New York Genome Center |
| Abecasis, Gonçalo | University of Michigan |
| Aguet, Francois | Broad Institute |
| Albert, Christine | Cedars Sinai |
| Almasy, Laura | Children's Hospital of Philadelphia, University of Pennsylvania |
| Alonso, Alvaro | Emory University |
| Ament, Seth | University of Maryland |
| Anderson, Peter | University of Washington |
| Anugu, Pramod | University of Mississippi |
| Applebaum-Bowden, Deborah | National Institutes of Health |
| Ardlie, Kristin | Broad Institute |
| Arking, Dan | Johns Hopkins University |
| Arnett, Donna K | University of Kentucky |
| Ashley-Koch, Allison | Duke University |
| Aslibekyan, Stella | University of Alabama |
| Assimes, Tim | Stanford University |
| Auer, Paul | University of Wisconsin Milwaukee |
| Avramopoulos, Dimitrios | Johns Hopkins University |
| Ayas, Najib | Providence Health Care |
| Balasubramanian, Adithya | Baylor College of Medicine Human Genome Sequencing Center |
| Barnard, John | Cleveland Clinic |
| Barnes, Kathleen | University of Colorado Anschutz Medical Campus |
| Barr, R. Graham | Columbia University |
| Barron-Casella, Emily | Johns Hopkins University |
| Barwick, Lucas | The Emmes Corporation |
| Beaty, Terri | Johns Hopkins University |
| Beck, Gerald | Cleveland Clinic |
| Becker, Diane | Johns Hopkins University |
| Becker, Lewis | Johns Hopkins University |
| Beer, Rebecca | National Heart, Lung, and Blood Institute, National Institutes of Health |
| Beitelshees, Amber | University of Maryland |
| Benjamin, Emelia | Boston University, Massachusetts General Hospital |
| Benos, Takis | University of Pittsburgh |
| Bezerra, Marcos | Fundação de Hematologia e Hemoterapia de Pernambuco - Hemope |
| Bielak, Larry | University of Michigan |
| Bis, Joshua | University of Washington |
| Blackwell, Thomas | University of Michigan |
| Blangero, John | University of Texas Rio Grande Valley School of Medicine |
| Boerwinkle, Eric | University of Texas Health at Houston |
| Bowden, Donald W. | Wake Forest Baptist Health |
| Bowler, Russell | National Jewish Health |
| Brody, Jennifer | University of Washington |
| Broeckel, Ulrich | Medical College of Wisconsin |
| Broome, Jai | University of Washington |
| Brown, Deborah | University of Texas Health at Houston |
| Bunting, Karen | New York Genome Center |
| Burchard, Esteban | University of California, San Francisco |
| Bustamante, Carlos | Stanford University |
| Buth, Erin | University of Washington |
| Cade, Brian | Brigham & Women's Hospital |
| Cardwell, Jonathan | University of Colorado at Denver |
| Carey, Vincent | Brigham & Women's Hospital |
| Carrier, Julie | University of Montreal |
| Carty, Cara | Washington State University |
| Casaburi, Richard | University of California, Los Angeles |
| Casas Romero, Juan P | Brigham & Women's Hospital |
| Casella, James | Johns Hopkins University |
| Castaldi, Peter | Brigham & Women's Hospital |
| Chaffin, Mark | Broad Institute |
| Chang, Christy | University of Maryland |
| Chang, Yi-Cheng | National Taiwan University |
| Chasman, Daniel | Brigham & Women's Hospital |
| Chavan, Sameer | University of Colorado at Denver |
| Chen, Bo-Juen | New York Genome Center |
| Chen, Wei-Min | University of Virginia |
| Chen, Yii-Der Ida | Lundquist Institute |
| Cho, Michael | Brigham & Women's Hospital |
| Choi, Seung Hoan | Broad Institute |
| Chuang, Lee-Ming | National Taiwan University |
| Chung, Mina | Cleveland Clinic |
| Chung, Ren-Hua | National Health Research Institute Taiwan |
| Clish, Clary | Broad Institute |
| Comhair, Suzy | Cleveland Clinic |
| Conomos, Matthew | University of Washington |
| Cornell, Elaine | University of Vermont |
| Correa, Adolfo | University of Mississippi |
| Crandall, Carolyn | University of California, Los Angeles |
| Crapo, James | National Jewish Health |
| Cupples, L. Adrienne | Boston University |
| Curran, Joanne | University of Texas Rio Grande Valley School of Medicine |
| Curtis, Jeffrey | University of Michigan |
| Custer, Brian | Vitalant Research Institute |
| Damcott, Coleen | University of Maryland |
| Darbar, Dawood | University of Illinois at Chicago |
| David, Sean | University of Chicago |
| Davis, Colleen | University of Washington |
| Daya, Michelle | University of Colorado at Denver |
| de Andrade, Mariza | Mayo Clinic |
| de las Fuentes, Lisa | Washington University in St Louis |
| de Vries, Paul | University of Texas Health at Houston |
| DeBaun, Michael | Vanderbilt University |
| Deka, Ranjan | University of Cincinnati |
| DeMeo, Dawn | Brigham & Women's Hospital |
| Devine, Scott | University of Maryland |
| Dinh, Huyen | Baylor College of Medicine Human Genome Sequencing Center |
| Doddapaneni, Harsha | Baylor College of Medicine Human Genome Sequencing Center |
| Duan, Qing | University of North Carolina |
| Dugan-Perez, Shannon | Baylor College of Medicine Human Genome Sequencing Center |
| Duggirala, Ravi | University of Texas Rio Grande Valley School of Medicine |
| Durda, Jon Peter | University of Vermont |
| Dutcher, Susan K. | Washington University in St Louis |
| Eaton, Charles | Brown University |
| Ekunwe, Lynette | University of Mississippi |
| El Boueiz, Adel | Harvard University |
| Ellinor, Patrick | Massachusetts General Hospital |
| Emery, Leslie | University of Washington |
| Erzurum, Serpil | Cleveland Clinic |
| Farber, Charles | University of Virginia |
| Farek, Jesse | Baylor College of Medicine Human Genome Sequencing Center |
| Fingerlin, Tasha | National Jewish Health |
| Flickinger, Matthew | University of Michigan |
| Fornage, Myriam | University of Texas Health at Houston |
| Franceschini, Nora | University of North Carolina |
| Frazar, Chris | University of Washington |
| Fu, Mao | University of Maryland |
| Fullerton, Stephanie M. | University of Washington |
| Fulton, Lucinda | Washington University in St Louis |
| Gabriel, Stacey | Broad Institute |
| Gan, Weiniu | National Heart, Lung, and Blood Institute, National Institutes of Health |
| Gao, Shanshan | University of Colorado at Denver |
| Gao, Yan | University of Mississippi |
| Gass, Margery | Fred Hutchinson Cancer Research Center |
| Geiger, Heather | New York Genome Center |
| Gelb, Bruce | Icahn School of Medicine at Mount Sinai |
| Geraci, Mark | University of Pittsburgh |
| Germer, Soren | New York Genome Center |
| Gerszten, Robert | Beth Israel Deaconess Medical Center |
| Ghosh, Auyon | Brigham & Women's Hospital |
| Gibbs, Richard | Baylor College of Medicine Human Genome Sequencing Center |
| Gignoux, Chris | Stanford University |
| Gladwin, Mark | University of Pittsburgh |
| Glahn, David | Boston Children's Hospital, Harvard Medical School |
| Gogarten, Stephanie | University of Washington |
| Gong, Da-Wei | University of Maryland |
| Goring, Harald | University of Texas Rio Grande Valley School of Medicine |
| Graw, Sharon | University of Colorado Anschutz Medical Campus |
| Gray, Kathryn J. | Mass General Brigham |
| Grine, Daniel | University of Colorado at Denver |
| Gross, Colin | University of Michigan |
| Gu, C. Charles | Washington University in St Louis |
| Guan, Yue | University of Maryland |
| Guo, Xiuqing | Lundquist Institute |
| Gupta, Namrata | Broad Institute |
| Haas, David M. | Indiana University |
| Haessler, Jeff | Fred Hutchinson Cancer Research Center |
| Hall, Michael | University of Mississippi |
| Han, Yi | Baylor College of Medicine Human Genome Sequencing Center |
| Hanly, Patrick | University of Calgary |
| Harris, Daniel | University of Maryland |
| Hawley, Nicola L. | Yale University |
| He, Jiang | Tulane University |
| Heavner, Ben | University of Washington |
| Heckbert, Susan | University of Washington |
| Hernandez, Ryan | University of California, San Francisco |
| Herrington, David | Wake Forest Baptist Health |
| Hersh, Craig | Brigham & Women's Hospital |
| Hidalgo, Bertha | University of Alabama |
| Hixson, James | University of Texas Health at Houston |
| Hobbs, Brian | Brigham & Women's Hospital |
| Hokanson, John | University of Colorado at Denver |
| Hong, Elliott | University of Maryland |
| Hoth, Karin | University of Iowa |
| Hsiung, Chao (Agnes) | National Health Research Institute Taiwan |
| Hu, Jianhong | Baylor College of Medicine Human Genome Sequencing Center |
| Hung, Yi-Jen | Tri-Service General Hospital National Defense Medical Center |
| Huston, Haley | Blood Works Northwest |
| Hwu, Chii Min | Taichung Veterans General Hospital Taiwan |
| Irvin, Marguerite Ryan | University of Alabama |
| Jackson, Rebecca | Oklahoma State University Medical Center |
| Jain, Deepti | University of Washington |
| Jaquish, Cashell | National Heart, Lung, and Blood Institute, National Institutes of Health |
| Johnsen, Jill | Blood Works Northwest |
| Johnson, Andrew | National Heart, Lung, and Blood Institute, National Institutes of Health |
| Johnson, Craig | University of Washington |
| Johnston, Rich | Emory University |
| Jones, Kimberly | Johns Hopkins University |
| Kang, Hyun Min | University of Michigan |
| Kaplan, Robert | Albert Einstein College of Medicine |
| Kardia, Sharon | University of Michigan |
| Kelly, Shannon | University of California, San Francisco |
| Kenny, Eimear | Icahn School of Medicine at Mount Sinai |
| Kessler, Michael | University of Maryland |
| Khan, Alyna | University of Washington |
| Khan, Ziad | Baylor College of Medicine Human Genome Sequencing Center |
| Kim, Wonji | Harvard University |
| Kimoff, John | McGill University |
| Kinney, Greg | University of Colorado at Denver |
| Konkle, Barbara | Blood Works Northwest |
| Kooperberg, Charles | Fred Hutchinson Cancer Research Center |
| Kramer, Holly | Loyola University |
| Lange, Christoph | Harvard School of Public Health |
| Lange, Ethan | University of Colorado at Denver |
| Lange, Leslie | University of Colorado at Denver |
| Laurie, Cathy | University of Washington |
| Laurie, Cecelia | University of Washington |
| LeBoff, Meryl | Brigham & Women's Hospital |
| Lee, Jiwon | Brigham & Women's Hospital |
| Lee, Sandra | Baylor College of Medicine Human Genome Sequencing Center |
| Lee, Wen-Jane | Taichung Veterans General Hospital Taiwan |
| LeFaive, Jonathon | University of Michigan |
| Levine, David | University of Washington |
| Levy, Dan | National Heart, Lung, and Blood Institute, National Institutes of Health |
| Lewis, Joshua | University of Maryland |
| Li, Xiaohui | Lundquist Institute |
| Li, Yun | University of North Carolina |
| Lin, Henry | Lundquist Institute |
| Lin, Honghuang | Boston University |
| Lin, Xihong | Harvard School of Public Health |
| Liu, Simin | Brown University |
| Liu, Yongmei | Duke University |
| Liu, Yu | Stanford University |
| Loos, Ruth J.F. | Icahn School of Medicine at Mount Sinai |
| Lubitz, Steven | Massachusetts General Hospital |
| Lunetta, Kathryn | Boston University |
| Luo, James | National Heart, Lung, and Blood Institute, National Institutes of Health |
| Magalang, Ulysses | Ohio State University |
| Mahaney, Michael | University of Texas Rio Grande Valley School of Medicine |
| Make, Barry | Johns Hopkins University |
| Manichaikul, Ani | University of Virginia |
| Manning, Alisa | Broad Institute, Harvard University, Massachusetts General Hospital |
| Manson, JoAnn | Brigham & Women's Hospital |
| Martin, Lisa | George Washington University |
| Marton, Melissa | New York Genome Center |
| Mathai, Susan | University of Colorado at Denver |
| Mathias, Rasika | Johns Hopkins University |
| May, Susanne | University of Washington |
| McArdle, Patrick | University of Maryland |
| McDonald, Merry-Lynn | University of Alabama |
| McFarland, Sean | Harvard University |
| McGarvey, Stephen | Brown University |
| McGoldrick, Daniel | University of Washington |
| McHugh, Caitlin | University of Washington |
| McNeil, Becky | RTI International |
| Mei, Hao | University of Mississippi |
| Meigs, James | Massachusetts General Hospital |
| Menon, Vipin | Baylor College of Medicine Human Genome Sequencing Center |
| Mestroni, Luisa | University of Colorado Anschutz Medical Campus |
| Metcalf, Ginger | Baylor College of Medicine Human Genome Sequencing Center |
| Meyers, Deborah A | University of Arizona |
| Mignot, Emmanuel | Stanford University |
| Mikulla, Julie | National Heart, Lung, and Blood Institute, National Institutes of Health |
| Min, Nancy | University of Mississippi |
| Minear, Mollie | National Institute of Child Health and Human Development, National Institutes of Health |
| Minster, Ryan L | University of Pittsburgh |
| Mitchell, Braxton D. | University of Maryland |
| Moll, Matt | Brigham & Women's Hospital |
| Momin, Zeineen | Baylor College of Medicine Human Genome Sequencing Center |
| Montasser, May E. | University of Maryland |
| Montgomery, Courtney | Oklahoma Medical Research Foundation |
| Muzny, Donna | Baylor College of Medicine Human Genome Sequencing Center |
| Mychaleckyj, Josyf C | University of Virginia |
| Nadkarni, Girish | Icahn School of Medicine at Mount Sinai |
| Naik, Rakhi | Johns Hopkins University |
| Naseri, Take | Ministry of Health, Government of Samoa |
| Natarajan, Pradeep | Broad Institute |
| Nekhai, Sergei | Howard University |
| Nelson, Sarah C. | University of Washington |
| Neltner, Bonnie | University of Colorado at Denver |
| Nessner, Caitlin | Baylor College of Medicine Human Genome Sequencing Center |
| Nickerson, Deborah | University of Washington |
| Nkechinyere, Osuji | Baylor College of Medicine Human Genome Sequencing Center |
| North, Kari | University of North Carolina |
| O'Connell, Jeff | University of Maryland |
| O'Connor, Tim | University of Maryland |
| Ochs-Balcom, Heather | University at Buffalo |
| Okwuonu, Geoffrey | Baylor College of Medicine Human Genome Sequencing Center |
| Pack, Allan | University of Pennsylvania |
| Paik, David T. | Stanford University |
| Palmer, Nicholette | Wake Forest Baptist Health |
| Pankow, James | University of Minnesota |
| Papanicolaou, George | National Heart, Lung, and Blood Institute, National Institutes of Health |
| Parker, Cora | RTI International |
| Peloso, Gina | Boston University |
| Peralta, Juan Manuel | University of Texas Rio Grande Valley School of Medicine |
| Perez, Marco | Stanford University |
| Perry, James | University of Maryland |
| Peters, Ulrike | Fred Hutchinson Cancer Research Center |
| Peyser, Patricia | University of Michigan |
| Phillips, Lawrence S | Emory University |
| Pleiness, Jacob | University of Michigan |
| Pollin, Toni | University of Maryland |
| Post, Wendy | Johns Hopkins University |
| Powers Becker, Julia | University of Colorado at Denver |
| Preethi Boorgula, Meher | University of Colorado at Denver |
| Preuss, Michael | Icahn School of Medicine at Mount Sinai |
| Psaty, Bruce | University of Washington |
| Qasba, Pankaj | National Heart, Lung, and Blood Institute, National Institutes of Health |
| Qiao, Dandi | Brigham & Women's Hospital |
| Qin, Zhaohui | Emory University |
| Rafaels, Nicholas | University of Colorado at Denver |
| Raffield, Laura | University of North Carolina |
| Rajendran, Mahitha | Baylor College of Medicine Human Genome Sequencing Center |
| Ramachandran, Vasan S. | Boston University |
| Rao, D.C. | Washington University in St Louis |
| Rasmussen-Torvik, Laura | Northwestern University |
| Ratan, Aakrosh | University of Virginia |
| Redline, Susan | Brigham & Women's Hospital |
| Reed, Robert | University of Maryland |
| Reeves, Catherine | New York Genome Center |
| Regan, Elizabeth | National Jewish Health |
| Reiner, Alex | Fred Hutchinson Cancer Research Center, University of Washington |
| Reupena, Muagututi‘a Sefuiva | Lutia I Puava Ae Mapu I Fagalele |
| Rice, Ken | University of Washington |
| Rich, Stephen | University of Virginia |
| Robillard, Rebecca | University of Ottawa |
| Robine, Nicolas | New York Genome Center |
| Roden, Dan | Vanderbilt University |
| Roselli, Carolina | Broad Institute |
| Rotter, Jerome | Lundquist Institute |
| Ruczinski, Ingo | Johns Hopkins University |
| Runnels, Alexi | New York Genome Center |
| Russell, Pamela | University of Colorado at Denver |
| Ruuska, Sarah | Blood Works Northwest |
| Ryan, Kathleen | University of Maryland |
| Sabino, Ester Cerdeira | Universidade de Sao Paulo |
| Saleheen, Danish | Columbia University |
| Salimi, Shabnam | University of Maryland |
| Salvi, Sejal | Baylor College of Medicine Human Genome Sequencing Center |
| Salzberg, Steven | Johns Hopkins University |
| Sandow, Kevin | Lundquist Institute |
| Sankaran, Vijay G. | Harvard University |
| Santibanez, Jireh | Baylor College of Medicine Human Genome Sequencing Center |
| Schwander, Karen | Washington University in St Louis |
| Schwartz, David | University of Colorado at Denver |
| Sciurba, Frank | University of Pittsburgh |
| Seidman, Christine | Harvard Medical School |
| Seidman, Jonathan | Harvard Medical School |
| Sériès, Frédéric | Université Laval |
| Sheehan, Vivien | Emory University |
| Sherman, Stephanie L. | Emory University |
| Shetty, Amol | University of Maryland |
| Shetty, Aniket | University of Colorado at Denver |
| Sheu, Wayne Hui-Heng | Taichung Veterans General Hospital Taiwan |
| Shoemaker, M. Benjamin | Vanderbilt University |
| Silver, Brian | UMass Memorial Medical Center |
| Silverman, Edwin | Brigham & Women's Hospital |
| Skomro, Robert | University of Saskatchewan |
| Smith, Albert Vernon | University of Michigan |
| Smith, Jennifer | University of Michigan |
| Smith, Josh | University of Washington |
| Smith, Nicholas | University of Washington |
| Smith, Tanja | New York Genome Center |
| Smoller, Sylvia | Albert Einstein College of Medicine |
| Snively, Beverly | Wake Forest Baptist Health |
| Snyder, Michael | Stanford University |
| Sofer, Tamar | Brigham & Women's Hospital |
| Sotoodehnia, Nona | University of Washington |
| Stilp, Adrienne M. | University of Washington |
| Storm, Garrett | University of Colorado at Denver |
| Streeten, Elizabeth | University of Maryland |
| Su, Jessica Lasky | Brigham & Women's Hospital |
| Sung, Yun Ju | Washington University in St Louis |
| Sylvia, Jody | Brigham & Women's Hospital |
| Szpiro, Adam | University of Washington |
| Taliun, Daniel | University of Michigan |
| Tang, Hua | Stanford University |
| Taub, Margaret | Johns Hopkins University |
| Taylor, Kent D. | Lundquist Institute |
| Taylor, Matthew | University of Colorado Anschutz Medical Campus |
| Taylor, Simeon | University of Maryland |
| Telen, Marilyn | Duke University |
| Thornton, Timothy A. | University of Washington |
| Threlkeld, Machiko | University of Washington |
| Tinker, Lesley | Fred Hutchinson Cancer Research Center |
| Tirschwell, David | University of Washington |
| Tishkoff, Sarah | University of Pennsylvania |
| Tiwari, Hemant | University of Alabama |
| Tong, Catherine | University of Washington |
| Tracy, Russell | University of Vermont |
| Tsai, Michael | University of Minnesota |
| Vaidya, Dhananjay | Johns Hopkins University |
| Van Den Berg, David | University of Southern California |
| VandeHaar, Peter | University of Michigan |
| Vrieze, Scott | University of Minnesota |
| Walker, Tarik | University of Colorado at Denver |
| Wallace, Robert | University of Iowa |
| Walts, Avram | University of Colorado at Denver |
| Wang, Fei | University of Washington |
| Wang, Heming | Brigham & Women's Hospital, Mass General Brigham |
| Wang, Jiongming | University of Michigan |
| Watson, Karol | University of California, Los Angeles |
| Watt, Jennifer | Baylor College of Medicine Human Genome Sequencing Center |
| Weeks, Daniel E. | University of Pittsburgh |
| Weinstock, Joshua | University of Michigan |
| Weir, Bruce | University of Washington |
| Weiss, Scott T | Brigham & Women's Hospital |
| Weng, Lu-Chen | Massachusetts General Hospital |
| Wessel, Jennifer | Indiana University |
| Willer, Cristen | University of Michigan |
| Williams, Kayleen | University of Washington |
| Williams, L. Keoki | Henry Ford Health System |
| Wilson, Carla | Brigham & Women's Hospital |
| Wilson, James | Beth Israel Deaconess Medical Center |
| Winterkorn, Lara | New York Genome Center |
| Wong, Quenna | University of Washington |
| Wu, Joseph | Stanford University |
| Xu, Huichun | University of Maryland |
| Yanek, Lisa | Johns Hopkins University |
| Yang, Ivana | University of Colorado at Denver |
| Yu, Ketian | University of Michigan |
| Zekavat, Seyedeh Maryam | Broad Institute |
| Zhang, Yingze | University of Pittsburgh |
| Zhao, Snow Xueyan | National Jewish Health |
| Zhao, Wei | University of Michigan |
| Zhu, Xiaofeng | Case Western Reserve University |
| Zody, Michael | New York Genome Center |
| Zoellner, Sebastian | University of Michigan |

**Supplementary Table 2:** All genome-wide significant ($P<5\times{10}^{-8})$ peaks from the recovery of GWAS signals for COPD meta-analysis.

| **MarkerName** | **Allele1** | **Freq1** | **OR** | **P-value** | **Direction** |
| --- | --- | --- | --- | --- | --- |
| 15:78817929:C:T | t | 0.61 | 0.78 | 1.33E-16 | --- |
| 15:78857986:C:G | c | 0.64 | 0.78 | 1.60E-16 | --- |
| 15:78867482:A:C | a | 0.63 | 0.78 | 1.91E-16 | --- |
| 15:78898723:C:T | t | 0.40 | 1.28 | 3.35E-16 | +++ |
| 15:78802869:C:T | t | 0.62 | 0.78 | 1.05E-15 | --- |
| 15:78900647:A:G | a | 0.65 | 0.78 | 1.48E-15 | --- |
| 15:78901113:C:T | t | 0.35 | 1.28 | 1.48E-15 | +++ |
| 15:78900701:T:G | t | 0.65 | 0.78 | 1.48E-15 | --- |
| 15:78900650:C:T | t | 0.35 | 1.28 | 1.48E-15 | +++ |
| 15:78816057:G:T | t | 0.36 | 1.28 | 1.67E-15 | +++ |
| 15:78899560:CAA:C | caa | 0.64 | 0.78 | 1.67E-15 | --- |
| 15:78899003:C:A | a | 0.35 | 1.28 | 1.77E-15 | +++ |
| 15:78898932:C:G | c | 0.65 | 0.78 | 2.14E-15 | --- |
| 15:78859605:AAAAAG:A | a | 0.36 | 1.28 | 2.20E-15 | +++ |
| 15:78850501:AT:A | a | 0.36 | 1.28 | 2.94E-15 | +++ |
| 15:78862064:C:T | t | 0.36 | 1.29 | 3.16E-15 | +++ |
| 15:78900908:C:T | t | 0.35 | 1.28 | 3.58E-15 | +++ |
| 15:78868636:G:A | a | 0.36 | 1.29 | 3.77E-15 | +++ |
| 15:78911181:T:C | t | 0.58 | 0.79 | 4.38E-15 | --- |
| 15:78865197:A:G | a | 0.65 | 0.78 | 6.00E-15 | --- |
| 15:78886198:C:T | t | 0.36 | 1.29 | 6.03E-15 | +++ |
| 15:78882925:G:A | a | 0.36 | 1.29 | 6.39E-15 | +++ |
| 15:78915370:CT:C | ct | 0.58 | 0.79 | 6.75E-15 | --- |
| 15:78866445:G:A | a | 0.35 | 1.28 | 7.05E-15 | +++ |
| 15:78906177:A:T | a | 0.64 | 0.78 | 7.39E-15 | --- |
| 15:78806023:T:C | t | 0.64 | 0.78 | 7.43E-15 | --- |
| 15:78862453:C:A | a | 0.36 | 1.28 | 7.43E-15 | +++ |
| 15:78886947:G:A | a | 0.36 | 1.29 | 7.45E-15 | +++ |
| 15:78813155:G:A | a | 0.36 | 1.28 | 8.64E-15 | +++ |
| 15:78873993:A:T | a | 0.64 | 0.78 | 9.22E-15 | --- |
| 15:78849034:T:C | t | 0.64 | 0.79 | 9.24E-15 | --- |
| 15:78851615:G:A | a | 0.36 | 1.27 | 1.30E-14 | +++ |
| 15:78857939:T:G | t | 0.64 | 0.78 | 1.66E-14 | --- |
| 15:78801394:A:C | a | 0.65 | 0.79 | 1.82E-14 | --- |
| 15:78912710:T:TGCGCGGGGCAGGGCGACGGGCA | t | 0.64 | 0.78 | 1.91E-14 | --- |
| 15:78878541:G:A | a | 0.36 | 1.27 | 2.01E-14 | +++ |
| 15:78877381:C:A | a | 0.36 | 1.27 | 2.20E-14 | +++ |
| 15:78894339:G:A | a | 0.35 | 1.27 | 2.50E-14 | +++ |
| 15:78911672:G:C | c | 0.36 | 1.27 | 2.78E-14 | +++ |
| 15:78814046:G:A | a | 0.63 | 0.79 | 3.72E-14 | --- |
| 15:78828086:G:T | t | 0.36 | 1.26 | 8.35E-14 | +++ |
| 15:78849779:C:T | t | 0.36 | 1.26 | 1.34E-13 | +++ |
| 15:78896129:A:G | a | 0.64 | 0.80 | 2.89E-13 | --- |
| 15:78826180:G:A | a | 0.64 | 0.80 | 1.10E-12 | --+ |
| 15:78915872:C:A | a | 0.38 | 1.24 | 3.43E-12 | +++ |
| 15:78915864:G:A | a | 0.44 | 1.23 | 5.41E-12 | +++ |
| 15:78914534:C:T | t | 0.38 | 1.23 | 1.32E-11 | +++ |
| 4:89866713:T:C | t | 0.57 | 0.83 | 2.66E-10 | --- |
| 4:89860830:C:T | t | 0.43 | 1.21 | 2.86E-10 | +++ |
| 4:89750361:A:G | a | 0.66 | 0.81 | 4.13E-10 | --? |
| 4:89869332:T:C | t | 0.55 | 0.83 | 4.19E-10 | --+ |
| 4:89870964:C:T | t | 0.45 | 1.21 | 4.70E-10 | ++- |
| 4:89869078:C:T | t | 0.44 | 1.20 | 6.10E-10 | ++- |
| 4:89873092:A:G | a | 0.56 | 0.83 | 8.89E-10 | --+ |
| 4:89869918:G:A | a | 0.44 | 1.20 | 8.89E-10 | ++- |
| 4:89872176:G:A | a | 0.44 | 1.20 | 1.15E-09 | ++- |
| 4:145463567:C:T | t | 0.38 | 0.83 | 2.12E-09 | --- |
| 4:145454374:C:T | t | 0.37 | 0.83 | 2.72E-09 | --- |
| 4:145449525:G:GAT | g | 0.63 | 1.20 | 3.51E-09 | +++ |
| 4:145437014:T:C | t | 0.43 | 1.19 | 3.70E-09 | +++ |
| 4:145465768:T:C | t | 0.63 | 1.20 | 3.73E-09 | +++ |
| 4:145462588:A:C | a | 0.63 | 1.20 | 3.73E-09 | +++ |
| 4:145458484:T:C | t | 0.63 | 1.20 | 3.73E-09 | +++ |
| 4:145460230:A:G | a | 0.63 | 1.20 | 3.73E-09 | +++ |
| 4:145455550:C:T | t | 0.37 | 0.83 | 3.73E-09 | --- |
| 4:145464885:A:G | a | 0.59 | 1.19 | 3.98E-09 | +++ |
| 4:145471245:A:T | a | 0.63 | 1.20 | 3.98E-09 | +++ |
| 4:145463521:G:A | a | 0.37 | 0.83 | 4.85E-09 | --- |
| 4:145463533:T:C | t | 0.63 | 1.20 | 4.85E-09 | +++ |
| 4:145460338:A:G | a | 0.63 | 1.20 | 4.85E-09 | +++ |
| 4:145465461:C:G | c | 0.63 | 1.20 | 4.85E-09 | +++ |
| 4:145434901:A:G | a | 0.62 | 1.20 | 4.98E-09 | +++ |
| 4:145454232:C:T | t | 0.37 | 0.84 | 5.07E-09 | --- |
| 4:145454964:T:C | t | 0.63 | 1.20 | 6.70E-09 | +++ |
| 4:145465491:T:G | t | 0.59 | 1.19 | 6.80E-09 | +++ |
| 4:145445779:A:T | a | 0.61 | 1.21 | 6.88E-09 | ++? |
| 4:145471255:CA:C | ca | 0.63 | 1.20 | 7.42E-09 | +++ |
| 4:145469968:G:T | t | 0.37 | 0.84 | 7.79E-09 | --- |
| 4:145474297:G:A | a | 0.37 | 0.84 | 7.79E-09 | --- |
| 4:145478662:C:T | t | 0.37 | 0.84 | 8.32E-09 | --- |
| 4:145473196:G:A | a | 0.38 | 0.84 | 1.10E-08 | --- |
| 4:89885714:T:C | t | 0.48 | 0.84 | 1.17E-08 | --- |
| 4:145436324:T:C | t | 0.63 | 1.19 | 1.27E-08 | +++ |
| 4:145436894:G:C | c | 0.37 | 0.84 | 1.27E-08 | --- |
| 15:78910258:C:T | t | 0.21 | 0.81 | 1.31E-08 | --- |
| 15:78910598:C:CT | ct | 0.22 | 0.82 | 1.46E-08 | --- |
| 4:145434744:A:G | a | 0.62 | 1.19 | 1.63E-08 | +++ |
| 15:78799060:ACT:A | a | 0.60 | 1.19 | 1.80E-08 | +++ |
| 19:41302706:C:T | t | 0.54 | 1.18 | 1.91E-08 | +++ |
| 4:145470657:T:A | a | 0.37 | 0.84 | 1.96E-08 | --- |
| 4:145478201:C:T | t | 0.37 | 0.84 | 1.96E-08 | --- |
| 4:145479880:T:C | t | 0.63 | 1.19 | 1.96E-08 | +++ |
| 4:145478049:C:G | c | 0.63 | 1.19 | 1.96E-08 | +++ |
| 4:145452783:C:T | t | 0.57 | 0.85 | 2.19E-08 | --- |
| 15:78899719:G:A | a | 0.77 | 1.22 | 2.25E-08 | +++ |
| 4:89884114:TG:T | t | 0.52 | 1.18 | 2.25E-08 | +++ |
| 4:89883979:C:T | t | 0.52 | 1.18 | 2.25E-08 | +++ |
| 15:78896547:G:A | a | 0.78 | 1.22 | 2.33E-08 | +++ |
| 15:78899213:T:C | t | 0.22 | 0.82 | 2.36E-08 | --- |
| 15:78912943:G:A | a | 0.21 | 0.82 | 2.49E-08 | --- |
| 15:78903987:T:C | t | 0.22 | 0.82 | 2.59E-08 | --- |
| 15:78888234:A:ACCCC | a | 0.78 | 1.22 | 2.75E-08 | +++ |
| 4:145463231:G:A | a | 0.38 | 0.85 | 2.75E-08 | --- |
| 4:145467212:A:T | a | 0.62 | 1.18 | 2.87E-08 | +++ |
| 15:78796104:C:T | t | 0.59 | 1.19 | 2.88E-08 | +++ |
| 15:78754000:A:G | a | 0.41 | 0.84 | 2.88E-08 | --- |
| 4:89924725:A:C | a | 0.55 | 1.19 | 3.06E-08 | ++? |
| 15:78765290:T:G | t | 0.41 | 0.84 | 3.08E-08 | --- |
| 4:89885086:G:T | t | 0.51 | 1.18 | 3.11E-08 | +++ |
| 4:145445117:A:G | a | 0.42 | 1.18 | 3.12E-08 | +++ |
| 15:78751961:C:T | t | 0.59 | 1.19 | 3.18E-08 | +++ |
| 15:78779510:T:A | a | 0.59 | 1.19 | 3.31E-08 | +++ |
| 15:78789223:G:A | a | 0.59 | 1.19 | 3.31E-08 | +++ |
| 4:89908381:GA:G | g | 0.33 | 0.84 | 3.31E-08 | --- |
| 15:78793921:C:T | t | 0.59 | 1.19 | 3.33E-08 | +++ |
| 15:78792398:T:C | t | 0.41 | 0.84 | 3.33E-08 | --- |
| 15:78747916:AAAAAAG:A | a | 0.59 | 1.19 | 3.35E-08 | +++ |
| 4:145479139:A:G | a | 0.59 | 1.18 | 3.37E-08 | +++ |
| 4:145480780:A:G | a | 0.59 | 1.18 | 3.37E-08 | +++ |
| 15:78845110:A:G | a | 0.79 | 1.22 | 3.51E-08 | +++ |
| 15:78837673:C:T | t | 0.21 | 0.82 | 3.58E-08 | --- |
| 15:78782095:C:T | t | 0.59 | 1.19 | 3.64E-08 | +++ |
| 15:78789488:C:T | t | 0.59 | 1.19 | 3.64E-08 | +++ |
| 4:145468791:G:A | a | 0.57 | 0.85 | 3.66E-08 | --- |
| 15:78767346:G:A | a | 0.59 | 1.19 | 3.67E-08 | +++ |
| 4:145445694:T:A | a | 0.57 | 0.85 | 3.75E-08 | --- |
| 4:145444039:A:C | a | 0.43 | 1.18 | 3.75E-08 | +++ |
| 4:145464074:T:A | a | 0.40 | 0.85 | 4.00E-08 | --- |
| 4:145462364:G:A | a | 0.40 | 0.85 | 4.00E-08 | --- |
| 4:89765661:G:C | c | 0.35 | 1.18 | 4.06E-08 | ++- |
| 15:78908565:C:T | t | 0.21 | 0.80 | 4.18E-08 | --? |
| 15:78769130:A:G | a | 0.41 | 0.84 | 4.30E-08 | --- |
| 4:89924434:GAA:G | g | 0.46 | 0.85 | 4.43E-08 | --- |
| 4:145425936:A:G | a | 0.64 | 1.19 | 4.57E-08 | ++- |
| 15:78766194:T:A | a | 0.59 | 1.18 | 4.73E-08 | +++ |
| 15:78767850:C:T | t | 0.59 | 1.18 | 4.73E-08 | +++ |
| 15:78868398:C:T | t | 0.25 | 0.83 | 4.79E-08 | --- |

**Supplementary Table 3: Number of samples and SNPs in each step of the analysis.** Taking the test between COPDGene EA array data and WGS data (Analysis 1 in the technical comparison) as an example, the number of samples and SNPs are recorded for each step. Throughout the table, the steps are as labeled in **Figure 1**.

|  | **COPDGene EA array data** | | **COPDGene EA WGS data** | |
| --- | --- | --- | --- | --- |
|  | **#Samples** | **#SNPs** | **#Samples** | **#SNPs** |
| Step 1: Original array and WGS data | 6,670 | 648,530 | 6,507 | 626,638,014 |
| Step 2: After Extraction of array SNPs from WGS | 6,670 | 648,530 | 6,507 | 630,947 |
| Step 3: After QC | 6,664 | 639,505 | 6,507 | 614,283 |
|  | **Merged #Samples** | | **Merged #SNPs** | |
| Step 5: After merging, input for 1^st^ imputation | 6,501 | | 557,746 | |
| Step 6: Input for 2^nd^ imputation | 6,501 | | 484,502 | |
| Step 7: After 2^nd^ imputation | 6,501 | | 47,109,431 | |
| Step 8: After all filtering | 6,501 | | 6,974,901 | |
